# The impact of testosterone on paraventricular nucleus gene expression in male and female spontaneously hypertensive rats

**DOI:** 10.1101/2025.02.20.639267

**Authors:** Alex Paterson, Su-Yi Loh, Shadi Kadijeh Gholami, Mark F Rogers, Dharmani Devi Murugan, Lam Sau-Kuen, Mohammad Rais Mustafa, Benjamin P Ott, Prusha Balaratnam, Andre S Mecawi, David Murphy, Charles C. T. Hindmarch

## Abstract

**Background:** Hypertension is a polygenic, complex disease that impacts men and women differently; whilst the incidence of high blood pressure (BP) is roughly equal over a lifetime, men typically are at higher risk of developing the disease earlier in life, before 50 years of age. There is adequate evidence that the brain is critical for the BP setpoint. The paraventricular nucleus (PVN) of the hypothalamus is an integrative structure that can influence not only neurohumoral responses to blood pressure changes, but also sympathetic drive. Here we manipulate the androgenic status of both male and female spontaneously hypertensive rats (SHRs) to determine how this changes gene expression within the PVN of these animals.

**Methods:** SHR (8-weeks old) were either sham-operated or orchiectomized, whereas all females were oophorectomized, half of which received 10mg testosterone propionate subcutaneously. Mean arterial pressure (MAP) and testosterone (T) were measured by carotid cannulation and ELISA respectively. Sequencing was performed on hand-punched PVN sections and subjected to robust bioinformatic analysis.

**Results:** In total, 6,571 differentially regulated genes (DRGs) are regulated in the PVN of male and female rats. High T (endogenous or replaced) correlates with higher MAP in both sexes. Orchidectomy-induced T depletion resulted in the significant regulation of 5,104 genes, involved in thousands of biological roles, including ones related to hormone and sex-hormone signalling. In the female SHR, testosterone replacement in oophorectomized animals induced the regulation of 1,727 genes, sharing many biological functions with those in the high T males. We validated key genes by qPCR to determine false discovery rate.

**Conclusions:** T status in hypertensive rats correlates with MAP, and consistent changes in PVN transcriptome

## Introduction

Primary hypertension is the result of an intractable mosaic of modifiable and un- modifiable risk factors that is responsible for approximately 9.4 million deaths per year; genetics and lifestyle conspire to change the blood pressure set point that ultimately results in early death. Whilst both men and women are susceptible to hypertension and share a lifelong risk for the disease, younger men are more likely than age-matched women to develop hypertension. In contrast, postmenopausal women are more likely to have high blood pressure than younger women. This implies a sex-dependent stratification of risk factors that may be dependent upon circulating sex hormones.

Hypertension is a multi-system, multi-organ complex disease that relies on an interplay of feedback loops that establish set points that respond to fluctuations in blood pressure (BP). There is evidence that brainstem and hypothalamic structures are associated with the development and maintenance of hypertension in both animal models and human subjects (*1*). Firstly, hypertensive patients present with higher levels of the sympathetic neurotransmitter, norepinephrine and this increase correlates well with an increase of sympathetic nerve traffic (*2, 3*). Secondly, these increases are detected in different clinical conditions characterised by an increase in blood pressure such as age related, pregnancy induced and white coat hypertensions (*4, 5*), demonstrating that the increase in sympathetic nerve activity is independent of the condition from which the hypertension arose.

Regardless of the underlying cause, there is unequivocal evidence that sympathetic nervous system (SNS) hyperactivity is involved both in the set-point change and maintenance of high BP (*1, 6–10*). SNS output from preganglionic neurons in the intermediolateral (IML) cell column results in a noradrenergic driven elevation of total peripheral resistance and cardiac output that drives an increase in blood pressure. It is not only the baroreceptive circuitry that is able to modulate sympathetic nerve activity (SNA). The paraventricular nucleus (PVN) is an integrative structure in the hypothalamus that can integrate a myriad of physiological signals and generate an appropriate response (*11–19*). When blood pressure drops, the PVN can respond at both neurohumoral levels through hormone release from the pituitary, and at sympathetic levels through descending projections from parvocellular neurones in the PVN to the brainstem, notably the rostroventrolateral medulla (RVLM), and intermediolateral cell column (IML) (*20, 21*). The PVN regulates changes in sympathetic nerve activity involved in the regulation of both arterial pressure and blood volume (*22–24*), and so is the master regulator of the blood pressure set-point by the brain.

Whilst global data shows no difference in incidence according to sex (26.6% of men, 26.1% of women), age-specific rates of hypertension show that the prevalence of hypertension is higher in men than in women under 50 years old whereas the converse is true after that age (*25*). Over 70% of postmenopausal (>50yo) women are hypertensive (*26*) which results from a combination of risk factors such as obesity, dietary life-style and importantly a change in circulating ovarian hormones (*27*) as well as an increase in circulating Angiotensin II (*28*), a potent vasoconstrictor. The observation that hypertension is less pronounced in premenopausal human females compared to age-matched males is mirrored in several rat models (*29–32*). The Spontaneously Hypertensive Rat (SHR)(*13, 33–35*) is a genetic model of hypertension that develops and maintains hypertension from about 6 weeks without experimental manipulation. The SHR correlates well with human hypertension especially in terms of SNA, for example blood pressure change in young animals is preceded by elevations SNA, and PVN excitatory drive to the RVLM appears to be heightened compared to the normotensive control strain, the Wistar Kyoto (WKY) rat (*36*).

Many studies have focused on estrogen and its associated cardioprotective effects as being responsible for this sex difference, blaming its post-menopausal reduction for the increase in blood pressure. Importantly, there are marked differences between the male and female SHR in terms of blood pressure, with male SHRs displaying higher BP than age-matched females (*37–41*). Whilst there is evidence for androgens accounting for these differences between the sexes, little has been done to understand the underlying mechanisms at play. There are also sex-specific differences in cardiovascular and metabolic hormones in the PVN (*42*). Manipulation of testosterone in both male and female SHR modifies BP, for example castrated male SHRs have lower BP than intact controls (*37–41*) whereas testosterone treatment of oophorectomized female SHRs elevates BP (*40*). The impact of testosterone was further demonstrated in a cohort of over 1800 pre- and post- menopausal women, as an increase in the ratio of testosterone to estradiol was found to be a negative predictor for developing metabolic syndrome, including elevated blood pressure, during the menopausal transition (*43*). Both estrogen and androgens are capable of driving gene transcription through their roles as transcription factors; bound estrogen and androgen receptors can translocate to the nucleus and act as transcription factors through the binding of target genes with either estrogen(*44*) or androgen(*45*) response elements.

Here we recapitulate a model of androgen manipulation first presented by Reckelhoff (*40*) who showed that in the SHR, testosterone exacerbates hypertension. Intact sham SHR males have higher testosterone and mean arterial pressure (MAP) than do oophorectomized SHR females. Castration of male SHR drops testosterone and MAP to female OVX levels, whereas testosterone treatment to oophorectomized females drives MAP to sham male levels. We performed exploratory RNAsequencing on the paraventricular nucleus of these animals and present here those differentially regulated genes in the PVN. We also draw specific attention to those transcripts that are directly regulated in the same direction by testosterone in both male and female SHRs.

## Materials and methods

### Animals

All procedures were carried out with the approval from Institutional Animal Care and Use Committee (IACUC), Universiti Malaya (ethic number: 2014-05-07/physio/R/NS). Adult, 8 week-old female and male Wistar-Kyoto (WKY) and Spontaneously Hypertensive (SHR) rats were obtained from Animal Experimental Unit (AEU), Universiti Malaya. The animals are kept in a well-maintained environment with standardised temperature (22±1°C), humidity (50±5°C) and 12:12 hours light-dark cycle. All animals had free access to standard rodent chow and tap water *ad libitum* throughout the experiment. Eight experimental groups were established: Sham- operated intact WKY males, orchiectomized WKY Males, Sham-operated intact SHR Males, orchiectomized SHR Males, oophorectomized WKY females, oophorectomized WKY females with testosterone treatment, oophorectomized SHR females, oophorectomized SHR Females with testosterone treatment.

Sham-operation, orchidectomies or oophorectomies were carried out at 8 weeks of age under ketamine: xylazine (80:8 mg/kg, intraperitoneal) anaesthesia. After 2 weeks of recovery period, pre-prepared 19-mm-length silastic tubing (0.062 in ID, 9.125 in OD; Dow Corning) containing 10mg testosterone propionate (Sigma-Aldrich, USA) was subcutaneously implanted at the back of the shoulder of female oophorectomized rats animals. The remaining control groups were implanted with empty silastic tubing. All silastic tubing was replaced after 3 weeks.

### Blood pressure measurement

After 6 weeks of treatment periods, animals (n = 6 per group) were subjected to carotid cannulation procedures under pentobarbital sodium (60 mg/kg, intraperitoneal) anaesthesia. In brief, a small incision was first made in the neck for tracheostomy and carotid artery cannulation. The carotid artery was identified with the visual aid of the vagus nerve and a cannula pre-filled with heparinized saline (0.5 IU/ml) was inserted. The direct blood pressures were monitored and measured via the cannula connected to the power lab system (ADInstruments). The 10-minutes recorded data were analysed using LabChart version 6 and average systolic (SBP) and diastolic (DBP) blood pressure were obtained. The mean arterial blood pressures (MAP) were calculated based on the equation DBP + [(SBP-DBP)/3].

### Tissue and blood collection

At 16 weeks of age, all animals were euthanized via decapitation (between 0800 and 1200). Trunk bloods were quickly collected in chilled heparinized blood collection tubes and centrifuged at 3,000rpm, 20mins at 4°C. Their brains were also removed from the skull and snap frozen on powdered dry ice. All plasma and frozen brains were stored at -80°C until further analysis.

### Measurement of testosterone

The measurement of testosterone levels in the blood plasma (n = 6 per group) were carried using enzyme-linked immunosorbent assay (ELISA) kit by Enzo Life Science, USA (Cat No. ADI-900-065) following the manufacturer’s protocols. Each assay was performed in triplicate and the lowest sensitivity limit of the assay was 5.67 pg/ml.

### RNA extraction for RNAseq

Tissues from the SHR comparisons only (n = 3 per group where 5 samples were pooled to obtain n = 1) were collected as discussed earlier. The frozen brain was first mounted onto Tissue-Tek OCT compound (Sakura Finetek, USA) on a chuck and placed in the cryostat (Shandon Cryotome FE and FSE Cryostats, Thermo Scientific). 60µm rostral-to-caudal sections were sliced and stained with Toluidine Blue (Sigma Aldrich, India). The sections were mapped using The Rat Brain: In Stereotaxic Coordinates, 6th edition and a 1-mm diameter sample corer was used to collect PVN, from un-stained tissues. QIAzol lysis reagent (Qiagen, Maryland, USA) was added into each sample and 5 samples were pooled to form an n = 1. The separations of RNA, DNA and protein phase were initiated by adding chloroform and centrifugation at 12,000xg, 15mins at 4°C. Upper layer was removed and RNA was precipitated with 1 volume of 70% (v/v) EtOH. The following RNA purification steps were carried out using Qiagen RNeasy Mini Kit (Qiagen, Maryland, USA) according to manufacturer’s protocol. The quality of the total RNA extracted were assessed using a Nanodrop (ThemoScientific NanoDrop 2000/2000c) and Agilent 2100 bioanalyzer instrument (Agilent Technologies, CA, USA; Agilent RNA 6000 Nano Kit). The concentrations of the RNA were quantified using Qubit RNA HS Assay Kit (ThermoFisher Scientific) following the manufacturer’s instructions on Qubit 2.0 Fluorometer (Invitrogen, Life technologies).

### RNAseq

Amplified cDNA libraries were prepared from isolated RNA samples and sequenced using the Illumina HiSeq 2500 Sequencer (Illumina Inc., USA) on HighOutput mode. Briefly, 1µg RNA from each pooled sample with RIN>8 and A260/A280: 2.0 was taken forward for sequencing. This was followed by the construction of Illumina libraries using ScriptSeq Complete Gold Kit (Illumina Inc., USA) that makes use of hybridisation to bead-bound rRNA probes to obtain rRNA depleted samples before applying unique barcode adapters. The libraries were assessed for their quality using a Qubit dsDNA High Sensitivity DNA Kit and their size determined by Agilent 2100 Bioanalyzer (Agilent Technologies, CA, USA; Agilent High Sensitivity DNA Kit). This followed with further enrichment and amplification of the libraries by qPCR using KAPA Biosystems Library Quantification Kit, then all samples were normalised to 4nM. Equal volumes of individual libraries (36 samples) were pooled and run on a MiSeq using MiSeq Reagent Kit v3 (Illumina) to validate the library clustering efficiency. The libraries were then re-pooled based on the MiSeq demultiplexing results and sequenced on a HiSeq 2500 sequencing platform (Illumina, San Diego, California, USA) and cBot with Ver 3 flow cells and sequencing reagents. Library reads of greater than 30 to 35 million were generated for each individual library. The data were then processed using RTA and CASAVA thus providing four sets of compressed FASTQ files per library. All raw reads were pre-processed for quality assessment, adaptor removal, quality trimming and size selection using the FASTQC toolkit to generate quality plots for all read libraries. We adopted a phred30 quality cutoff (99.9% base call accuracy).

### RNAsequencing analysis pipeline

RNAseq alignment and analysis was performed in house using bespoke pipelines run on our high-power computer (named ‘Hydra’), a Dell PowerEdgeR820 48 core computer equipped with 512GB RAM. Our pipeline makes use of bash, R, and Python scripting to accept RNAseq pre-trimmed data as input, before ultimately producing output tables of differentially expressed transcripts. Read alignment is performed using Tophat (*46*), optimized for the rat genome; ENSEMBL Rn6 annotations are used to determine the distribution of known intron lengths and adjust the range of acceptable intron lengths to account for roughly 99.9% of known introns (size range = 12-270,000 nucleotides). We use HTSeq (*47*) to generate read counts, using the ENSEMBL Rn6 annotations for reference. In order to determine those genes that are differentially expressed (DE) in these brain regions between SHR or WKY (n=3 per group), our pipeline makes use of edgeR (*48*) and DESeq2 from the R Bioconductor package. These DE predictions allow us to establish high-confidence predictions that have low p-values. We used gprofiler (*49*) to organize our differentially regulated genes into functionally and/or biologically relevant groups split by either gene ontology (GO) terms relevant to Biological Process (BP), Molecular Function (MF), Cellular Component (CC), Kyoto Encyclopaedia of Genes and Genomes (KEGG), Reactome (REACT), Wiki Pathways (WP), Transfac (TF), MIRNA (miRTarBase), CORUM protein complexes (CORUM), Human Protein Atlas (HPA) and Human Phenotype Ontology (HP).

### Candidate Selection

Data was organized using a spreadsheet and is filtered according to the edgeR p- value with a 0.05 cut-off applied. EdgeR significant genes from the Sham vs ORX data set were cross-referenced with the OVX vs OVXT dataset for each tissue. Then, genes that were under an arbitrarily selected threshold of 50 reads (post normalization), were omitted from further processing in order to prevent false predictions of large fold changes. From this list, genes were ranked, highest to lowest, by fold change between the experimental groups. This gave a final list of differentially regulated genes that were potentially the result of high testosterone. Candidates for further validation were selected from both the up and down regulated genes based on literature reviews of their potential involvement in neurocardiovascular regulation

### RNA extraction and cDNA synthesis for PCR

Tissues (n = 6 per group) were collected and RNA were extracted using the similar methods as described earlier. 200-300ng of total RNA was used to synthesize cDNA using Quantitect reverse transcription kit (Qiagen, Maryland, USA) according to the manufacturer’s manuals. The mRNA levels in the tissues were assessed using QuantiNova SYBR Green PCR Kit (Qiagen, Germany) on the StepOne Plus Real-Time PCR Systems (Applied Biosystems). All primers were designed by using NCBI PrimerBLAST tool and synthesized by using Integrated DNA Technologies (Table 1). All qPCR assay were carried out in duplicate and the relative quantification of gene expression were obtained based on the 2^-ΔΔCt^ method. Because commonly used housekeeping genes can fluctuate according to tissue, we firstly assessed each condition for housekeeper gene CT value differences across each condition and report no significant difference in the expression of Rpl19, Gapdh or Actb according to condition or strain of animal. RPL19 were selected as the reference gene in order to normalized the gene expression levels between samples.

### Statistical analysis

With the exception of the RNAsequencing analysis, which was performed in R, all data analysis were performed using Graphpad Prism (Graphpad Software) and were expressed as Mean ± Standard error of Mean (SEM).

## Results

### Testosterone and mean arterial pressure in male WKY and SHR rats

In both WKY and SHR male animals, orchidectomy attenuated circulating testosterone compared to shams (**Figure 1A**). In addition, the basal sham testosterone of the SHR males (∼3ng/mL) was substantially higher than in the WKY males (∼2ng/mL). Mean arterial blood pressure (MAP) in both WKY and SHR correlated with the drop in testosterone seen in the orchiectomised animals (**Figure 1B/C**).

**Figure 1.**
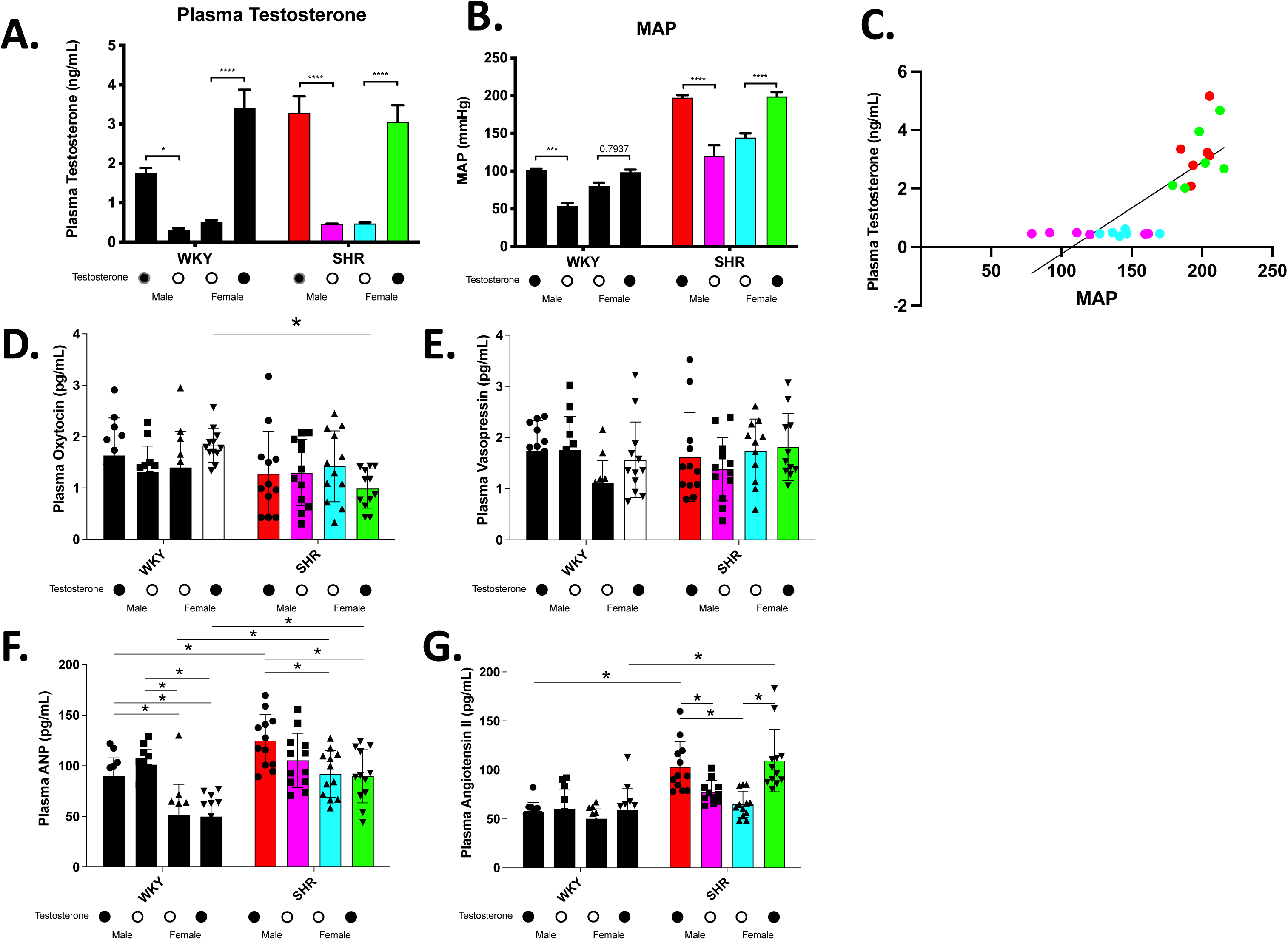
Manipulation of testosterone correlates with blood pressure changes in hypertensive rats: Plasma testosterone levels (**A**) and MAP (**B**) in intact or orchiectomized Male, and, intact or oophorectomized with testosterone repletion Female Wistar Kyoto (WKY) and Spontaneously Hypertensive (SHR) rats. Intact males in both strains have a higher testosterone level and a corresponding higher MAP than do intact females. When male WKY and SHR are orchiectomized, plasma testosterone and MAP drop to the level of intact females (no significant difference). When female animals from both strains are given testosterone, the WKY do not exhibit a significant change in MAP compared to intact female WKY, but in the SHR, MAP is comparable to intact males. **(C)** blood pressure significantly correlates with testosterone.gra In these same groups, we find no significant difference in the SHR of **(D)** oxytocin, or **(E)** vasopressin, however differences in the SHR and WKY are noticed in **(F)** atrial natriuretic peptide (ANP), **(G)** and angiotensin II. (\**p*<0.05, \*\**p*<0.01, \*\*\**p*<0.001, \*\*\*\**p*<0.0001)

### Testosterone and mean arterial pressure in female WKY and SHR rats

In both oophorectomized WKY and SHR rats, testosterone levels were low and comparable with orchiectomized male rats (**Figure 1A**). Testosterone supplementation significantly elevated circulating testosterone in both strains. The MAP of female WKY oophorectomized animals that received testosterone was not significantly different to the oophorectomized controls, however, testosterone supplemented female SHR MAP was significantly higher than oophorectomized controls (**Figure 1B**).

### Hormone profile of male and female WKY and SHR rats following androgen induction

We profiled the circulating levels of Vasopressin (AVP), Oxytocin (OXT), Angiotensin II (ANGII), and Atrial Natriuretic Peptide (ANP) to determine if androgen modulation in normotensive or hypertensive animals was responsible for changes in these cardiovascular peptides. No significant differences in OXT or AVP levels were identified in either the WKY or the SHR strains (**Figure 1 D/E**), or as a consequence of androgen modulation within strains. ANP levels were significantly higher in male SHR sham animals compared to WKY shams, and female animals with or without testosterone drive, however ANP was not modulated by androgen manipulation in these animals (**Figure 1F**).

ANGII was significantly higher in SHR sham males compared to WKY, and was significantly attenuated by orchidectomy in the SHRs, but not WKYs (**Figure 1G**). Androgen supplementation in the female SHR resulted in a significant elevation of ANGII that was comparable to sham male SHR levels.

### RNAseq data and target validation

#### Sham male SHR vs. oophorectomized female SHR

When the molecular footprint of the PVN was assessed using RNA sequencing, 6,571 genes were identified as being differentially regulated between male sham SHR compared and oophorectomized female SHR (2,949 Down, 3,622 Up; **Figure 2A; Supplemental Figure S1).** When we passed these list genes through functional analysis, we revealed 2,136 enriched terms (**Supplemental Figure S2**), including 14 terms related to the term ‘hormone’ (**Figure 2B, Supplemental Figure S3**), including the term ‘response to hormone’ (GO:BP; GO:0009725), a term that included 179 genes; ‘response to steroid hormone’ (GO:BP; GO:48545) which involved 67 genes, and; Estrogen signaling (WikiPathways = WP; WP1279) which involved 34 genes).

**Figure 2.**
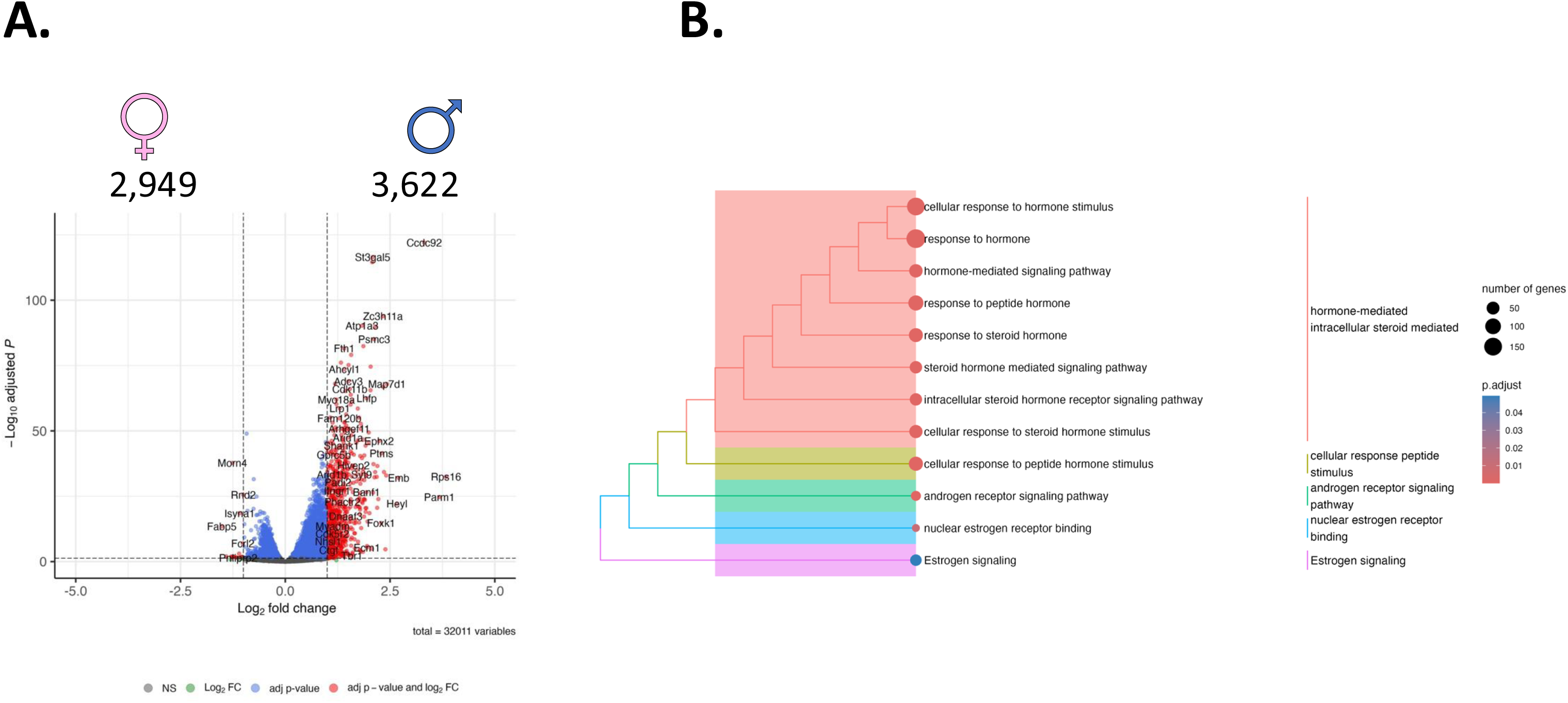
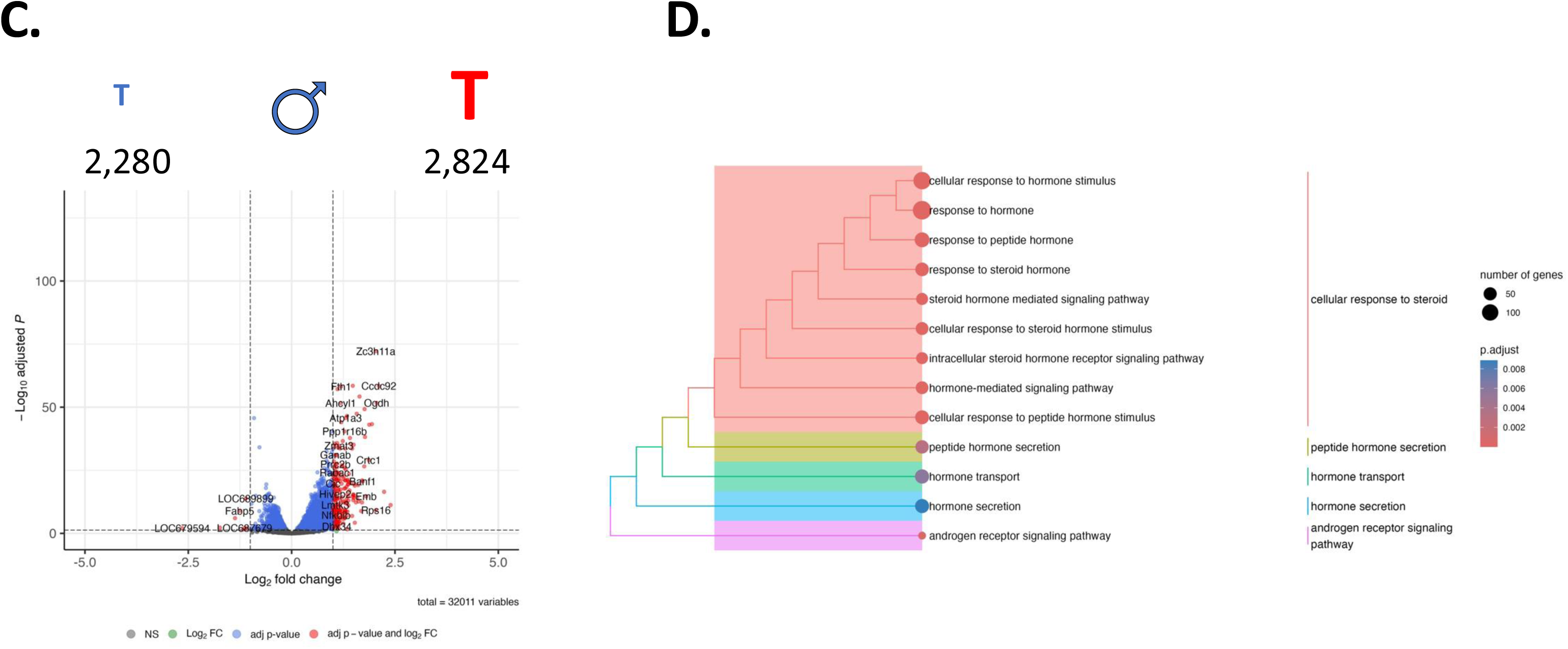
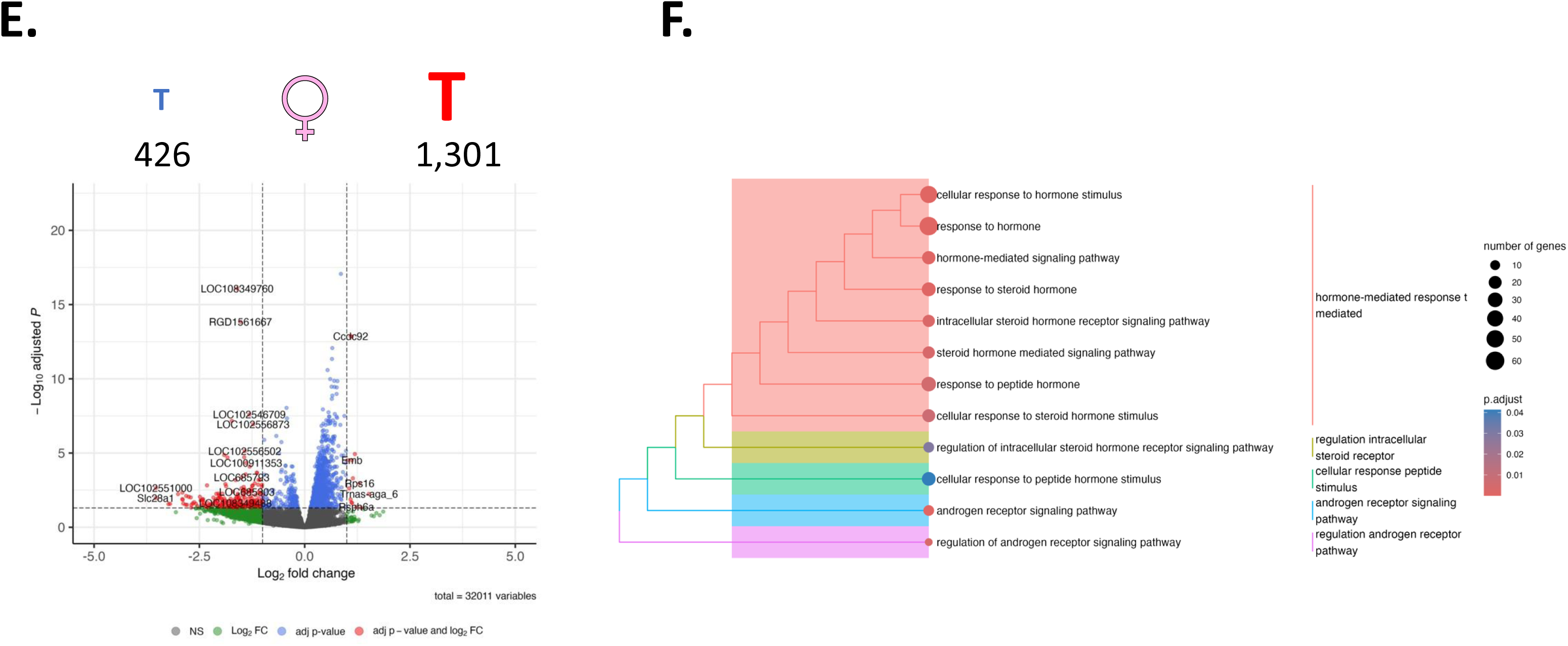
RNA sequencing of male and female hypertensive rats reveals differential gene expression profiles in the paraventricular nucleus which are consistent with androgen status: We report differential gene expression profiles that exist between; sham male and oophorectomized female spontaneously hypertensive rats (SHR; **A**); sham and orchiectomized male SHR’s (**C**); and oophorectomized and oophorectomized + testosterone female SHR’s (**E**). In each comparison, we report enriched gene functions that exist (**B, C, D**).

#### Sham male SHR vs. orchiectomised male SHR

When we compared transcriptomes between orchiectomized and intact males revealed 5,104 differentially regulated genes (2,280 Down, 2,824 Up; **Figure 2C; Supplemental Figure 4**). When these genes were parsed through gprofiler, we revealed 1,818 functional groups **(Supplemental Figure 5**), including 13 terms that were related to the term ‘hormone’ (**Figure 2D, Supplemental Figure 6**), that included term ‘hormone’, including the term ‘response to hormone’ (GO:BP; GO:0009725), a term that included 148 genes; and ‘androgen receptor signaling pathway’ (GO:BP; GO:0030521) that relates to 18 genes.

#### Oophorectomized female SHR vs. oophorectomized female SHR + testosterone

In the comparison between oophorectomized female SHR and oophorectomized female SHR supplemental with testosterone, 1,727 genes were differentially regulated (426 Down, 1,301 Up; **Figure 2E; Supplemental Figure 7).** G-profiler assessment of this set of genes revealed 960 functional groups (**Supplemental Figure 8**), including 12 that relate to the term ‘hormone’ (**Figure 2F, Supplemental Figure 9**) including ‘regulation of androgen receptor signaling pathway’ (GO:BP; GO:0030432), involving 8 genes, ‘androgen receptor signaling pathway’ (GO:BP; GO:0030521), involving 12 genes.

We wanted to validate targets from this dataset to establish our false discovery rate. We therefore used qPCR to validate the expression of 12 genes prioritized from our RNAseq data based on their regulation in response to testosterone (**Figure 3**); Each of these genes is upregulated in states of high testosterone; e.g. intact males, and females with exogenous testosterone. These genes are also enriched in the male PVN compared to the female PVN. We also validated both oxytocin (*11*) transcript, and heteronuclear transcript (OXT and hnOXT). We also validated the expression of epoxide hydrolase 2 (Ephx2) as we recognize this gene to be a marker of hypertension in various tissues in SHR compared to WKY (*unpublished observation).* This validation gives some confidence that the genes we report are not within the false discovery rate.

**Figure 3.**
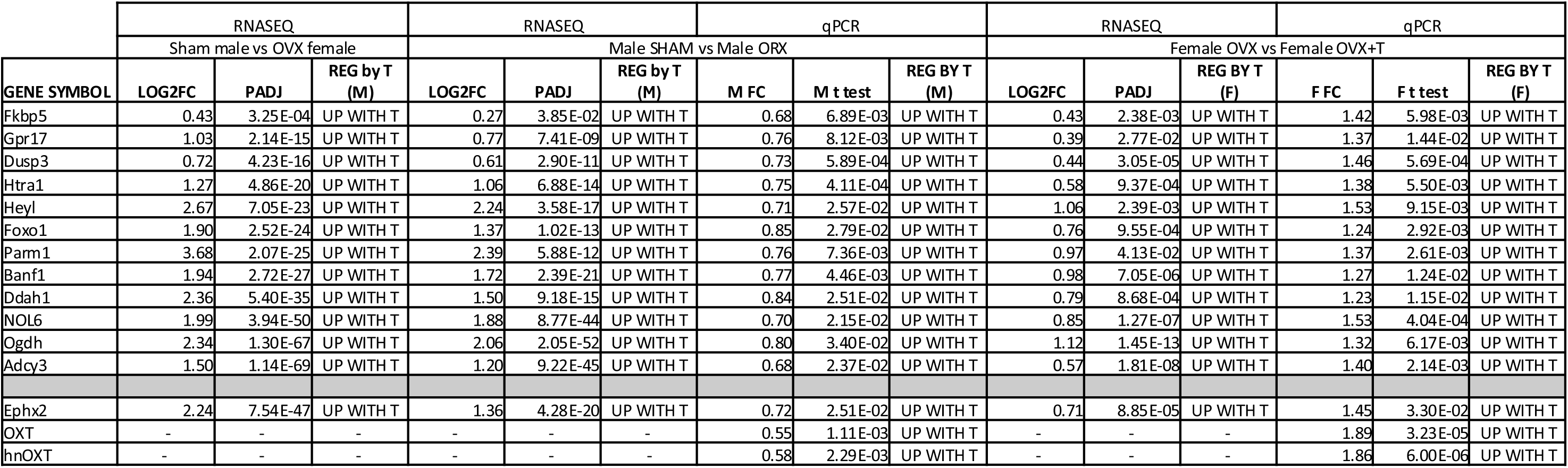
Genes that were identified in the RNA sequencing data as upregulated by testosterone in each comparison were validated by qPCR: We used quantitative (q)PCR to understand our false discovery rate. Here, we validated 12 genes that were up regulated in the sham male vs oophorectomized female data and validated the expression of these genes in the experiments where testosterone (T) was modulated; Male sham vs male orchidectomy (ORX), and, female oophorectomized (OXT) vs. female oophorectomized + testosterone. PADJ = adjusted p-value following multiple test correction; T = testosterone, FC = Fold Change; M = male; F = female.

## Discussion

The SHR rat is a genetically programmed model of hypertension whereby animals spontaneously develop hypertension at about 4-6 weeks of age without intervention or environmental manipulation (*50*). The SHR is a favoured model for human hypertension because the cause of the elevated blood pressure is polygenic in nature and results in elevations in both RAAS (*51*) and SNA (*52*). In addition, younger male rats have a higher blood pressure than age-matched females, consistent with the human condition whereby men are more likely to be diagnosed with hypertension earlier in life. Because menopause in humans causes blood pressure in females to increase beyond that of males, a role of androgens and oestrogens in hypertension has been suggested (*53*). In this study, we created a model of androgen manipulation in the SHR and have described the transcriptional profile of the PVN - a key central region responsible for modulating sympathetic tone.

Our model was successful in recapitulating previous models of testosterone manipulation. MAP for sham males was consistently higher than that for sham females, whereas androgen depletion by orchidectomy dropped blood pressure in the males, in contrast, oophorectomy followed by testosterone supplementation in females elevated blood pressure to sham male levels. Our model demonstrates that the manipulation of a steroid hormone is sufficient to dictate the cardiovascular endpoint in a genetic model of hypertension.

We wanted to better understand the role of the PVN in this transition as we know that the PVN is not only capable of regulating blood pressure at the neurohumoral level, but also at the sympathetic level via direct connection to the RVLM. We identified 6,571 genes that are differentially regulated in the PVN of male SHRs following orchidectomy, and 1,727 differentially regulated genes in the female SHR PVN supplemented with testosterone. We additionally find 6,571 genes that are differentially regulated in the PVN as a consequence of sex. We have previously reported that there are significant sex-specific differences in the regulation of cardiovascular and metabolic hormones exist in in the PVN are sex-dependent (*42*).

One of the regulated genes we validated was epoxide hydrolayse 2 (Ephx2). Several studies have noted the significant increase in Ephx2 in the SHR when compared to its normotensive control the WKY (*54–56*). Indeed, many of our own datasets support this finding, to the degree that Ephx2 has become a putative marker for the SHR in several central brain regions (*unpublished*). The data we present here supports this significant difference between the SHR and WKY strains and further shows a profile of reduction in the absence of testosterone. Again, this is consistent with blood pressure reduction. Ephx2 encodes a protein that is one of the soluble epoxide hydrolases (sEH), responsible for converting epoxyeicosatrienoic acids (EETs) to dihydroxyeicosatrienoic acids (DHETs) via hydrolase activities (*57*). As EETs participate in a wide range of biological processes including cardiac contractility, inflammation and regulation of vascular tone, the blockage of sEH has become an attractive prospect for treating cardiovascular and inflammatory diseases (*58, 59*). One of the few examples investigating the direct relationship between testosterone and Ephx2 in mice showed that a Ephx2 reduction resulted in lower testosterone levels in C57BL6 Ephx2^(-/-)^ null mice, which were found to otherwise develop normally, and not display any obvious symptoms of disease or organ malfunction. However, they did have significantly lower levels of circulating plasma testosterone, as well as a reduced sperm count and testicular size, secondary sex characteristics that are resultant of testosterone actions (*57*).

Whilst we did not observe a significant change in *Oxt* or *Avp* transcript expression in our sequencing data, we wanted to independently examine the expression of these RNAs using qPCR because the PVN is one of the central sites for the synthesis, manufacture, and transport of these hormones to the periphery, and to central structures. We did not find a significant change in the expression of *Avp* in the PVN, however we did notice an increase in *Oxt*-derived transcripts (both nuclear and heteronuclear RNA) in line with testosterone levels; *Oxt* is upregulated in male shams relative to castrated animals and upregulated in females receiving testosterone compared to OVX shams. In psychiatric disorders, in which the PVN is a relevant structure, *Oxt* expression is under the control of the androgen receptor (AR) in its role as a transcription factor. Treatment of neuroblastoma cells with testosterone reduced *Oxt* mRNA levels significantly, the result of AR binding to an Androgen Receptor Element (*60*). While this is contrary to our finding here, we note that it demonstrates the influence of AR on OXT expression. We also note that OXT has an established role in decreasing blood pressure peripherally that is manifest through central mechanisms (*61*).

We also identified 12 genes that were prioritized using our RNAseq data, and which were successfully validated by qPCR in independent cohorts of WKY and SHR animals. Whilst these gene lists are exploratory, the regulation patterns are in line with genes that are upregulated in physiology where testosterone is high; sham males, and OVX females with testosterone treatment. Two of the genes we validated are relevant to known literature in the PVN (*Adcy3* and *Fkbp5*), both of which are involved in stress related functions within the PVN which is part of the hypothalamo-pituitary axis (HPA) that responds to glucocorticoid feedback in the stress response. *Adcy3* (adenylate cyclase 3), is a gene encoding an enzyme that catalyses the synthesis of the second messenger cyclic AMP. *Adcy3* is implicated in obesity (*62*), and *Adcy3* null mice experience hyperphagia and obesity (Wang et al 2009). Previous studies of microarray and qPCR data in the WKY have shown *Adcy3* to be up-regulated in arterial vessels in response to an adrenocorticotrophin hormone (ACTH) induced hypertension paradigm (*63*). KBP5 is involved in the stress response and epigenetic regulation, through interactions with glucocorticoid receptor activity and environmental stressors (49). *Fkbp5* deletion results in a dampened acute stress response, commensurate with an increase in glucocorticoid sensitivity, whereas overexpression of *Fkbp5* in the PVN results in hypothalamic pituitary axis over-activation (*64*). Emerging research is surrounding the HPA playing a critical role in stress response and the differential expression of FKBP5, is indicative of hypertension mediation through stress pathways, alluding to interactions between stress, epigenetics, and hypertension (51). FKBP5 has significant roles in the cellular process occurring in the periphery and brain contributing to several stress-related disorders (51). The other genes we have validated are orphan genes within the PVN, whose function(s) are unknown. These 12 genes are all up-regulated in the PVN of hypertensive animals with high testosterone, irrespective of sex. Overall, the differential expression of these genes emphasizes the multifaceted nature of testosterone-driven hypertension. Each gene presents unique pathways and mechanisms involved in hypertension and these genes highlight the potential therapeutic target genes of interest.

We also organized our datasets into enriched functional groups so that we could gain insights into the overarching roles these genes. We have highlighted those enriched functions linked to wildcard terms like ‘hormone’, ‘androgens’ or ‘oestrogens’, and determined overlapping functional groups that are shared in the PVN of hypertensive animals with differential androgen states. Common themes of these functions include ‘regulation of androgen receptor signaling pathway’ (GO:BP; GO:0030432), ‘androgen receptor signaling pathway’ (GO:BP; GO:0030521), steroid mediated signaling pathway (GO:BP; GO:0043401), and cellular response to steroid hormone stimulus (GO:BP; GO:0043401). Taken together, the genes that are differentially regulated in the PVN of rats in response to high, or low testosterone environments assemble into common functional pathways that are involved in response to steroid hormones like androgens.

Whilst our data does not explore this, we are cognisant that when androgens are bound to their receptor, this complex can translocate to the nucleus and acts as a transcription factor (*44, 45, 65*). We hypothesise that a high androgen environment, irrespective of the sex of the animal drives increases in MAP, which are managed in part by the regulation of genes in the PVN which may be under the regulatory control of bound androgen receptors.

## Limitations

Animal models are always flawed because they are simultaneously an artificial manifestation of symptoms that mirror human health and disease, but also because they present confounding factors unrelated to those symptoms. For example, we have already reported transcriptome specific sex differences in a related tissue, the supraoptic nucleus; rats and mice manifest different gene expression, but through the same pathways (*66*). A positive aspect of the SHR is that it requires no pharmacological treatment to elevate blood pressure, and that they display differences between sexes. The first issue with our experiment arises with the contrasting surgical procedures between the sexes and the potential for confounding factors; an oophorectomy is a far more invasive operation than an orchidectomy. However, it should be noted that guidelines support similar post-operative care between the surgeries (*67*). Furthermore, the assumption of sham and oophorectomy as being baselines for assessing gene expression under “control” conditions is not necessarily valid. For this reason, each sex was an isolated comparison and our attempts to draw a comparison between the sexes is made after the fact. We also concede that the inclusion of a sham-operated female would have provided better understanding of age-matched cycling SHRs. This strategy, however, presents potential issues with gene expression influences that arise from oestrus (*68, 69*), so this group was not assessed. Here the focus is on removing the possibly confounding role of oestrus in gene expression within these brain regions and therefore allowing the comparison between testosterone and a lack thereof to be made. Our RNAseq analysis utilised an n=3 per group; such low replication can call into question systems level analysis, due to the possibility of a high false discovery rate (FDR) (*70*). However, we have used qPCR to validate target genes and have focussed on discussion of functional terms that describe biological plausibility within our system, e.g. those functions involved in hormone signalling. Lastly, animals were euthanised by way of cranial strike followed by decapitation. This was performed in order to minimise the presence of potentially confounding gene expression signatures that result from pharmacological anaesthesia. A recent paper demonstrated this by use of *in situ* Fos staining to detect activation of a host anaesthesia-activated neuroendocrine neurons specifically within the hypothalamus (*71*).

## Conclusions

We have presented here data from the paraventricular nucleus of male or female spontaneously hypertensive rats exposed to different levels of circulating testosterone; male rats were either under baseline (high) testosterone levels, or attenuated levels because of castration, whereas females were either under basal (low) testosterone levels or exposed to artificial elevations in testosterone. We demonstrate that testosterone, mean arterial blood pressure and plasma angiotensin II levels change in line with testosterone levels irrespective of the sex of the animal. RNA sequencing revealed thousands of targets, of which a majority were under the control of testosterone acting within the role of a transcription factor. Our data will be useful to members of the scientific community interested in the regulation of sympathetic tone, blood pressure control, and testosterone induced cardiovascular changes.

### Perspectives and Significance

Although men and women have a comparable lifetime risk of hypertension, this risk is differentially stratified according to age, with younger men, and post-menopausal women being more likely to have higher blood pressure. Instead of focussing on the cardioprotective impact of oestrogen in hypertension, we are focused here on understanding the cardiovascular risk factor of androgens, specifically testosterone. Through manipulating testosterone in both males (through castration), and females (through testosterone treatment) in hypertensive animals, we have tried to describe the molecular environment of the paraventricular nucleus of the hypothalamus, so that we can better understand how this integrative structure contributes to cardiosympathetic tone, and neurohumoral response to androgen manipulation.

## DECLARATIONS

### Authors’ contributions

All authors read and approved the final manuscript. <u>Study conception and design</u>: MRM, ASM, DM, CCTH, <u>Animal husbandry and surgery</u>: SYL, KG, DDM, LSK, <u>Molecular biology</u>: AP, SYL, KG, DDM, <u>Hormone analysis</u>: ASM, <u>Bioinformatics and analysis</u>: AP, MR, CCTH, Manuscript preparation and writing: AP, DM, ASM, CCTH

## Acknowledgements

we acknowledge the contribution of Ms Chitra Devi in animal husbandry

## Ethics

All procedures were carried out with the approval from Institutional Animal Care and Use Committee (IACUC), Universiti Malaya (ethic number: 2014-05-07/physio/R/NS).

## References

1. G. Grassi, Role of the sympathetic nervous system in human hypertension. J Hypertens 16, 1979–1987 (1998).

2. W. J. Louis, A. E. Doyle, S. Anavekar, Plasma norepinephrine levels in essential hypertension. N Engl J Med 288, 599–601 (1973).

3. W. J. Louis, A. E. Doyle, S. N. Anavekar, Plasma noradrenaline concentration and blood pressure in essential hypertension, phaeochromocytoma and depression. Clin Sci Mol Med Suppl 2, 239s–242s (1975).

4. J. A. Benmoussa, M. Clarke, D. Bloomfield, White Coat Hyperglycemia: The Forgotten Syndrome. J Clin Med Res 8, 567–568 (2016).

5. K. H. Lampinen et al., Increased plasma norepinephrine levels in previously pre-eclamptic women. J Hum Hypertens 28, 269–273 (2014).

6. S. J. Mann, Neurogenic essential hypertension revisited: the case for increased clinical and research attention. Am J Hypertens 16, 881–888 (2003).

7. M. Esler, D. Kaye, Sympathetic nervous system activation in essential hypertension, cardiac failure and psychosomatic heart disease. J Cardiovasc Pharmacol 35, S1–7 (2000).

8. T. E. Lohmeier, J. R. Lohmeier, S. Warren, P. J. May, J. T. Cunningham, Sustained activation of the central baroreceptor pathway in angiotensin hypertension. Hypertension 39, 550–556 (2002).

9. M. J. Petersson et al., Increased cardiac sympathetic drive in renovascular hypertension. J Hypertens 20, 1181–1187 (2002).

10. K. Sakata et al., Potentiated sympathetic nervous and renin-angiotensin systems reduce nonlinear correlation between sympathetic activity and blood pressure in conscious spontaneously hypertensive rats. Circulation 106, 620–625 (2002).

11. M. Lozić et al., Overexpression of oxytocin receptors in the hypothalamic PVN increases baroreceptor reflex sensitivity and buffers BP variability in conscious rats. Br J Pharmacol 171, 4385–4398 (2014).

12. S. G. V. Dutra et al., Physiological and Transcriptomic Changes in the Hypothalamic-Neurohypophysial System after 24 h of Furosemide- Induced Sodium Depletion. Neuroendocrinology 111, 70–86 (2021).

13. A. Martin et al., Transcriptome Analysis Reveals Downregulation of Urocortin Expression in the Hypothalamo-Neurohypophysial System of Spontaneously Hypertensive Rats. Front Physiol 11, 599507 (2020).

14. G. G. Hazell et al., G protein-coupled receptors in the hypothalamic paraventricular and supraoptic nuclei--serpentine gateways to neuroendocrine homeostasis. Front Neuroendocrinol 33, 45–66 (2012).

15. C. C. T. Hindmarch, D. Murphy, The transcriptome and the hypothalamo- neurohypophyseal system. Endocr Dev 17, 1–10 (2010).

16. C. Hindmarch et al., The transcriptome of the rat hypothalamic- neurohypophyseal system is highly strain-dependent. J Neuroendocrinol 19, 1009–1012 (2007).

17. J. Qiu et al., Transcription factor expression in the hypothalamo- neurohypophyseal system of the dehydrated rat: upregulation of gonadotrophin inducible transcription factor 1 mRNA is mediated by cAMP-dependent protein kinase A. J Neurosci 27, 2196–2203 (2007).

18. M. T. Ghorbel et al., Microarray screening of suppression subtractive hybridization-PCR cDNA libraries identifies novel RNAs regulated by dehydration in the rat supraoptic nucleus. Physiol Genomics 24, 163–172 (2006).

19. C. Hindmarch, S. Yao, G. Beighton, J. Paton, D. Murphy, A comprehensive description of the transcriptome of the hypothalamoneurohypophyseal system in euhydrated and dehydrated rats. Proc Natl Acad Sci U S A 103, 1609–1614 (2006).

20. L. W. Swanson, P. E. Sawchenko, Paraventricular nucleus: a site for the integration of neuroendocrine and autonomic mechanisms. Neuroendocrinology 31, 410–417 (1980).

21. P. E. Sawchenko, L. W. Swanson, The organization of noradrenergic pathways from the brainstem to the paraventricular and supraoptic nuclei in the rat. Brain Res 257, 275–325 (1982).

22. Z. Yang, J. H. Coote, The influence of the paraventricular nucleus on baroreceptor dependent caudal ventrolateral medullary neurones of the rat. Pflugers Arch 438, 47–52 (1999).

23. S. G. Hardy, Hypothalamic projections to cardiovascular centers of the medulla. Brain Res 894, 233–240 (2001).

24. J. H. Coote, A role for the paraventricular nucleus of the hypothalamus in the autonomic control of heart and kidney. Exp Physiol 90, 169–173 (2005).

25. P. M. Kearney et al., Global burden of hypertension: analysis of worldwide data. Lancet 365, 217–223 (2005).

26. K. Sandberg, H. Ji, Sex differences in primary hypertension. Biol Sex Differ 3, 7 (2012).

27. C. Maric-Bilkan, E. L. Gilbert, M. J. Ryan, Impact of ovarian function on cardiovascular health in women: focus on hypertension. Int J Womens Health 6, 131–139 (2014).

28. S. S. Signorelli et al., Duration of menopause and behavior of malondialdehyde, lipids, lipoproteins and carotid wall artery intima- media thickness. Maturitas 39, 39–42 (2001).

29. M. Abdelbary et al., Necrosis Contributes to the Development of Hypertension in Male, but Not Female, Spontaneously Hypertensive Rats. Hypertension 74, 1524–1531 (2019).

30. J. C. Sullivan, K. Bhatia, T. Yamamoto, A. A. Elmarakby, Angiotensin (1-7) receptor antagonism equalizes angiotensin II-induced hypertension in male and female spontaneously hypertensive rats. Hypertension 56, 658–666 (2010).

31. L. A. Fortepiani, J. F. Reckelhoff, Role of oxidative stress in the sex differences in blood pressure in spontaneously hypertensive rats. J Hypertens 23, 801–805 (2005).

32. S. H. Lee et al., Sex-related differences in the intratubular renin- angiotensin system in two-kidney, one-clip hypertensive rats. Am J Physiol Renal Physiol 317, F670–F682 (2019).

33. K. Okamoto, K. Aoki, Development of a strain of spontaneously hypertensive rats. Jpn Circ J 27, 282–293 (1963).

34. H. Waki et al., Excessive leukotriene B4 in nucleus tractus solitarii is prohypertensive in spontaneously hypertensive rats. Hypertension 61, 194–201 (2013).

35. C. C. Hindmarch et al., The transcriptome of the medullary area postrema: the thirsty rat, the hungry rat and the hypertensive rat. Exp Physiol 96, 495–504 (2011).

36. A. M. Allen, Inhibition of the hypothalamic paraventricular nucleus in spontaneously hypertensive rats dramatically reduces sympathetic vasomotor tone. Hypertension 39, 275–280 (2002).

37. J. F. Reckelhoff, Sex steroids, cardiovascular disease, and hypertension: unanswered questions and some speculations. Hypertension 45, 170–174 (2005).

38. J. F. Reckelhoff, Gender differences in the regulation of blood pressure. Hypertension 37, 1199–1208 (2001).

39. J. F. Reckelhoff, H. Zhang, K. Srivastava, J. P. Granger, Gender differences in hypertension in spontaneously hypertensive rats: role of androgens and androgen receptor. Hypertension 34, 920–923 (1999).

40. J. F. Reckelhoff, H. Zhang, J. P. Granger, Testosterone exacerbates hypertension and reduces pressure-natriuresis in male spontaneously hypertensive rats. Hypertension 31, 435–439 (1998).

41. D. S. Martin, S. Biltoft, R. Redetzke, E. Vogel, Castration reduces blood pressure and autonomic venous tone in male spontaneously hypertensive rats. J Hypertens 23, 2229–2236 (2005).

42. S. P. Loewen et al., Sex-specific differences in cardiovascular and metabolic hormones with integrated signalling in the paraventricular nucleus of the hypothalamus. Exp Physiol 102, 1373–1379 (2017).

43. J. I. Torréns et al., Relative androgen excess during the menopausal transition predicts incident metabolic syndrome in midlife women: study of Women’s Health Across the Nation. Menopause 16, 257–264 (2009).

44. K. Ikeda, K. Horie-Inoue, S. Inoue, Identification of estrogen-responsive genes based on the DNA binding properties of estrogen receptors using high-throughput sequencing technology. Acta Pharmacol Sin 36, 24–31 (2015).

45. Q. Wang et al., A hierarchical network of transcription factors governs androgen receptor-dependent prostate cancer growth. Mol Cell 27, 380–392 (2007).

46. C. Trapnell, L. Pachter, S. L. Salzberg, TopHat: discovering splice junctions with RNA-Seq. Bioinformatics 25, 1105–1111 (2009).

47. S. Anders, P. T. Pyl, W. Huber, HTSeq—a Python framework to work with high-throughput sequencing data. Bioinformatics 31, 166–169 (2015).

48. M. D. Robinson, D. J. McCarthy, G. K. Smyth, edgeR: a Bioconductor package for differential expression analysis of digital gene expression data. Bioinformatics 26, 139–140 (2010).

49. U. Raudvere et al., g:Profiler: a web server for functional enrichment analysis and conversions of gene lists (2019 update). Nucleic Acids Res 47, W191–W198 (2019).

50. J. G. Dickhout, R. M. Lee, Blood pressure and heart rate development in young spontaneously hypertensive rats. Am J Physiol 274, H794–800 (1998).

51. K. Shiono, H. Sokabe, Renin-angiotensin system in spontaneously hypertensive rats. Am J Physiol 231, 1295–1299 (1976).

52. W. V. Judy et al., Sympathetic nerve activity: role in regulation of blood pressure in the spontaenously hypertensive rat. Circ Res 38, 21–29 (1976).

53. C. Moretti, G. Lanzolla, M. Moretti, L. Gnessi, E. Carmina, Androgens and Hypertension in Men and Women: a Unifying View. Curr Hypertens Rep 19, 44 (2017).

54. T. R. Harris, B. D. Hammock, Soluble epoxide hydrolase: gene structure, expression and deletion. Gene 526, 61–74 (2013).

55. D. Ai et al., Soluble epoxide hydrolase plays an essential role in angiotensin II-induced cardiac hypertrophy. Proc Natl Acad Sci U S A 106, 564–569 (2009).

56. J. M. Seubert et al., Differential renal gene expression in prehypertensive and hypertensive spontaneously hypertensive rats. Am J Physiol Renal Physiol 289, F552–561 (2005).

57. A. Luria et al., Alteration in plasma testosterone levels in male mice lacking soluble epoxide hydrolase. Am J Physiol Endocrinol Metab 297, E375–383 (2009).

58. N. Chiamvimonvat, C. M. Ho, H. J. Tsai, B. D. Hammock, The soluble epoxide hydrolase as a pharmaceutical target for hypertension. J Cardiovasc Pharmacol 50, 225–237 (2007).

59. K. Node et al., Anti-inflammatory properties of cytochrome P450 epoxygenase-derived eicosanoids. Science 285, 1276–1279 (1999).

60. D. Dai et al., Direct Involvement of Androgen Receptor in Oxytocin Gene Expression: Possible Relevance for Mood Disorders. Neuropsychopharmacology 42, 2064–2071 (2017).

61. M. Petersson, P. Alster, T. Lundeberg, K. Uvnäs-Moberg, Oxytocin causes a long-term decrease of blood pressure in female and male rats. Physiol Behav 60, 1311–1315 (1996).

62. H. Cao, X. Chen, Y. Yang, D. R. Storm, Disruption of type 3 adenylyl cyclase expression in the hypothalamus leads to obesity. Integr Obes Diabetes 2, 225–228 (2016).

63. T. H. Grayson et al., Vascular microarray profiling in two models of hypertension identifies caveolin-1, Rgs2 and Rgs5 as antihypertensive targets. BMC Genomics 8, 404 (2007).

64. A. S. Häusl et al., The co-chaperone Fkbp5 shapes the acute stress response in the paraventricular nucleus of the hypothalamus of male mice. Mol Psychiatry 26, 3060–3076 (2021).

65. S. Wang et al., Target analysis by integration of transcriptome and ChIP- seq data with BETA. Nat Protoc 8, 2502–2515 (2013).

66. L. Stewart et al., Hypothalamic transcriptome plasticity in two rodent species reveals divergent differential gene expression but conserved pathways. J Neuroendocrinol 23, 177–185 (2011).

67. A. I. Idris, Ovariectomy/orchidectomy in rodents. Methods Mol Biol 816, 545–551 (2012).

68. C. T. Minson, J. R. Halliwill, T. M. Young, M. J. Joyner, Influence of the menstrual cycle on sympathetic activity, baroreflex sensitivity, and vascular transduction in young women. Circulation 101, 862–868 (2000).

69. J. R. Carter, Q. Fu, C. T. Minson, M. J. Joyner, Ovarian cycle and sympathoexcitation in premenopausal women. Hypertension 61, 395–399 (2013).

70. T. F. Khang, C. Y. Lau, Getting the most out of RNA-seq data analysis. PeerJ 3, e1360 (2015).

71. L. F. Jiang-Xie et al., A Common Neuroendocrine Substrate for Diverse General Anesthetics and Sleep. Neuron 102, 1053–1065.e1054 (2019).

